# POISE: Spectral Inference of Parent-of-Origin Effects in Unlabeled Genomic Data

**DOI:** 10.64898/2026.06.10.731310

**Authors:** Ilha Hwang, Austin Talbot, S. Taylor Head, Cristina Trevino, Thomas S. Wingo, Alex Kotlar

## Abstract

**Motivation:** Parent of Origin Effects (POEs), where the effect of an an allele on a phenotype differs based on maternal or paternal inheritance implicated in growth, metabolism, and neurodevelopment. Traditional tests for POEs require family data to determine parental origins of transmitted alleles. Given that such studies are expensive and time consuming compared to genome-wide association studies (GWAS), tests that function absent inheritance information are highly desirable. We develop a method, based on community detection from machine learning, that infers POEs via a spectral decomposition, obtains confidence intervals via a non-parametric bootstrap, and safeguards against confounding by non POE sources of variation. We refer to our method as Parent of Origin Inference via Spectral Estimation (POISE).

**Results:** We demonstrate that POISE is well-calibrated under both Gaussian and heavy-tailed noise in simulation studies, with improved robustness to true POEs compared to existing covariance-based tests. POISE provides per-trait effect estimates with bias-corrected bootstrap confidence intervals and incorporates an information-theoretic minimum detectable effect size that filters unreliable estimates, conferring robustness to covariance-deflating variance QTL. We then apply POISE to GWAS data from the UK Biobank using BMI, LDL cholesterol, and HDL cholesterol. POISE recovers established POE loci and identifies 134 additional variants at genes implicated in lipid metabolism, immune regulation, and growth.

**Availability and implementation:** The code for this method in Python is available at https://github.com/bystrogenomics/POISE.

## 1. Introduction

A parent-of-origin effect (POE) is a discrepancy in the phenotypic impact of an allele depending on whether it is inherited from the mother or the father [Lawson et al., 2013]. A well-characterized mechanism underlying POEs is genomic imprinting, where genes are expressed from only one parental allele [Reik and Walter, 2001]. Imprinted genes play essential roles in growth, placentation, and neurodevelopment [McGrath and Solter, 1984, Surani et al., 1984, Barton et al., 1984], and their disregulation is implicated in a range of human disorders [Li et al., 1993, Lalande, 1996], such as Prader–Willi and Beckwith–Wiedemann syndromes [Nicholls et al., 1989, Weksberg et al., 1993]. As such, POEs have been proposed as a contributor to the “missing heritability” problem, the large gap between the phenotypic variance explained by genome-wide association studies (GWAS) and total estimated heritability [Guilmatre and Sharp, 2012].

Despite their importance, POEs remain difficult to detect at scale. Direct detection requires parental genotypes or trio data [Huxtable et al., 2000], allowing methods such as PREMIM/EMIM [Howey et al., 2015] to compare the two inheritance groups. However these data are scarce in large biobank cohorts. In the first UK Biobank cohort, for example, fewer than 5,000 of over 400,000 genotyped participants have directly available parental genotypes [Hofmeister et al., 2022], limiting scalability. To address this gap, variance-based methods have been developed that can detect POEs from GWAS of unrelated individuals, without requiring parental genotypes. These methods depend on mean differences between the groups inflating the heterozygote variances [Head et al., 2023]. Hoggart et al. [2014] introduced a univariate variance test for this signal, and POIROT extended it to the multivariate setting via a two-sample covariance test [Head et al., 2023].

While POIROT represents an important advance, it has several practical limitations. POIROT returns a single *p*-value per variant, testing whether the multivariate covariance structure differs between heterozygotes and homozygotes across a set of traits. However, it does not estimate effect sizes, much less quantify uncertainty in these estimates. Furthermore, the p-value used in POIROT is a general-purpose two-sample test and does not match the structure of an additive POE. As a result, it can falsely conflate a variance quantitative trait locus [Charcosset and Gallais, 1996], or vQTL, as a POE.

In this work, we introduce POISE (Parent of Origin Inference via Spectral Estimation), a method that addresses these limitations. Given a set of phenotypes and genotypes from unrelated individuals, POISE estimates the per-trait POE magnitudes at each locus, allowing researchers to determine which phenotypes are impacted by the POE. POISE accomplishes this by modeling the heterozygous population as a mixture and estimating the difference in covariance structure between heterozygotes and homozygotes to recover the POE coefficients. In addition to point estimates, POISE provides:

1. **Per-trait confidence intervals**. Using a bias-corrected and accelerated bootstrap procedure [Efron and Tibshirani, 1994, Efron, 1987], POISE produces confidence intervals for each trait-specific POE estimate, enabling uncertainty quantification that is essential for interpreting effect sizes.
2. **A sensitive covariance-based significance test**. This test can either be permutation-based or any two-sampled test.
3. **A principled reliability threshold**. POISE includes an effect-size criterion, derived from the statistical properties of the estimation problem, that distinguishes reliably estimated POEs from noise. This threshold depends on the minor allele frequency and sample size at each variant and acts as a safeguard: only effects large enough to be accurately estimated are reported, improving robustness to assumption violations.

## 2. Methods

### 2.1. Parent of Origin Effect and Notation

We first define notation following the conventions of Hoggart et al. [2014] and Head et al. [2023]. Consider a reference allele *A* and alternative allele *B*, defining the reference allele as the more frequent allele in the population. We observe *n*_*AA*_ homozygous individuals with an *AA* genotype and *n*_*AB*_ heterozygous individuals with the genotype *AB*. We exclude homozygotes of the alternative allele from analysis, as the heterozygous population is the critical population for effect estimates. Each individual also has *d* observed continuous phenotypes. The simplest model of the phenotypes for the *i*-th reference homozygote is

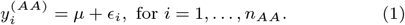

Here *µ* = (*µ*_1_, …, *µ*_*d*_)^*T*^ is the mean of the phenotypes and *ϵ*_*i*_ is the additive measurement noise. We assume that 𝔼[*ϵ*_*i*_] = 0 for identifiability of *µ*, and 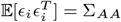.

We now model the heterozygous individuals. A standard GWAS would represent the heterozygote phenotype as

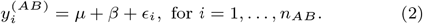

The parameter *β* represents the additive effect of the alternative allele on each phenotype, ignoring potential POEs. To incorporate a POE, we instead model the heterozygote phenotype as

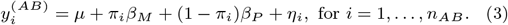

Here, *β*_*M*_ and *β*_*P*_ represent the maternal and paternal effects respectively and *π*_*i*_ indicates whether the allele came from the mother or father. In this model, the POE is Δ = *β*_*M*_ − *β*_*P*_ representing the difference between the effect of maternal and paternal inheritance. Furthermore, we assume that the covariance structure is shared across genotype groups: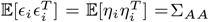, where *η*_*i*_ is the noise for heterozygotes.

### 2.2. The Basic POE Estimation Algorithm

With trio data one can separate the two inheriatance groups and compare their mean phenotypes, as shown on the left of Figure 1. This additive difference, Δ = *β*_*M*_ − *β*_*P*_, is the POE. Our method recovers Δ from heterozygotes even without these labels from the “bulge” indutroduced by the additive difference. We first describe the estimator in a simplified setting where phenotypes are assumed to be uncorrelated (Section 2.2.1), and then extend it to general case (Section 2.2.2).

**Fig. 1:**
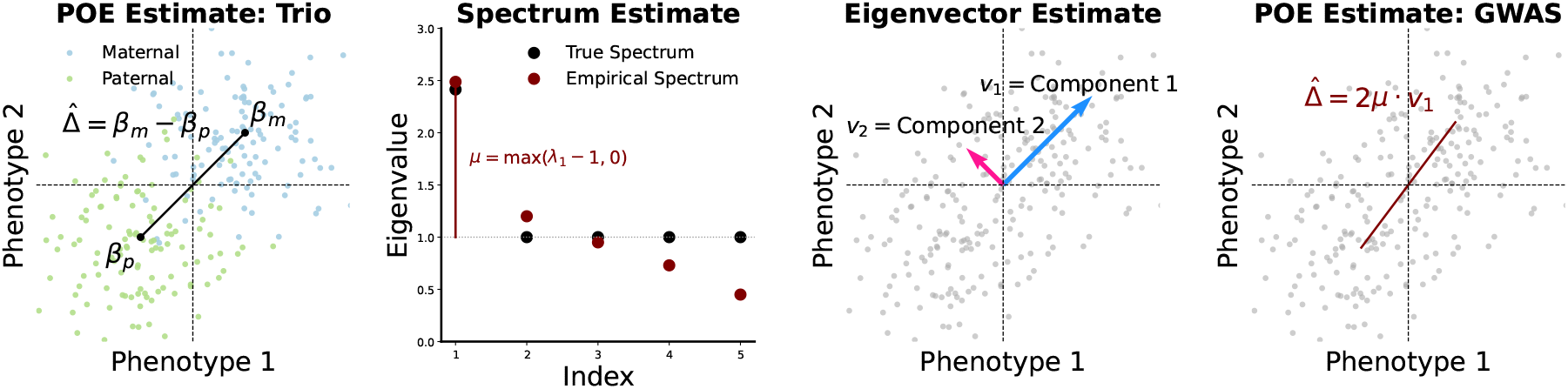
Illustrating the POE estimator with uncorrelated phenotypes (Σ_*AA*_ = *I*, two synthetic phenotypes). **(A)** The pooled, mean-centered heterozygous population as it is observed in practice: each point is one individual, and inheritance is unknown. The covariance ellipse (black) is noticeably elongated along one axis, hinting at a hidden structure. **(B)** The same data, but now colored by the (normally unobserved) inheritance pattern—red for individuals who inherited the alternative allele maternally, blue for paternally. The large circles mark the two group means, and the black arrow between them is the POE vector Δ = *β*_*M*_ − *β*_*P*_. In a trio study, Δ could be estimated directly as this difference in means. **(C)** Eigendecomposition of the empirical covariance from panel (A), requiring no inheritance information. The arrows show the two eigenvectors, each originating from the sample centroid (the origin after mean-centering) and scaled by the square root of its eigenvalue. The leading eigenvector (solid red, *λ*_1_ *>* 1) aligns with the POE direction; the second eigenvector (dashed blue, *λ*_2_ ≈ 1) is orthogonal and reflects noise only. The excess *λ*_1_ − 1 encodes the POE magnitude. **(D)** The per-phenotype POE estimate 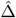 recovered via equation (7) (red bars), compared to the true Δ (dark bars).

#### 2.2.1. Simple Case: Uncorrelated Phenotypes

Assume that the phenotype covariance is the identity matrix 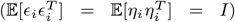. After subtracting the mean of the heterozygous population, the mean-centered phenotype of heterozygous individual *i* is

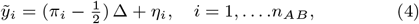

When we compute the covariance of the pooled heterozygous population without observing *π*_*i*_, there are two sources of variation: (i) the phenotypic noise *η*_*i*_, which is present regardless of any POE, and (ii) the separation between the two hidden inheritance groups. The law of total variance gives

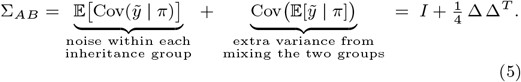

The first term is simply the phenotypic noise covariance *I*. The second term arises because the two group means (+Δ*/*2 and −Δ*/*2) are equally probable, producing a between-group variance of 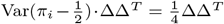. This rank-one “bump” is the POE signal we detect.

Since 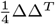 is rank one, it perturbs the identity in exactly one direction, the direction of Δ itself [Vershynin, 2018]. The eigenvalues of Σ_*AB*_ are 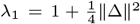 along the direction of Δ, and *λ*_2_ = *· · ·* = *λ*_*d*_ = 1 in all orthogonal directions, as shown in the second panel of Figure 1. That is, the POE “stretches” the covariance along the direction separating the two hidden groups (Component 1 of the third panel), and the amount of stretching encodes ∥Δ∥.

To estimate these parameters, we compute the empirical covariance of the mean-centered heterozygous population,

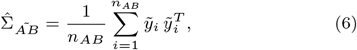

and extract its leading eigenvalue 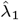 and eigenvector 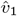. Inverting the eigenvalue relation gives 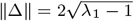, resulting in a POE estimate of

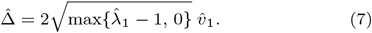

The max*{·*, 0*}* ensures that when 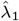 falls below 1 due to sampling noise or assumption violations, the estimate is zero rather than undefined. Each entry of 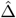 gives the estimated POE on the corresponding phenotype: large entries identify phenotypes that are strongly affected by parent of origin, while entries near zero indicate phenotypes that are not. The final estimate is visualized in the last panel of Figure 1.

#### 2.2.2. General Case: Correlated Phenotypes

In practice, phenotypes are correlated and have unequal variances (Σ_*AA*_≠ *I*). This is the typical situation in biobanks: metabolic traits co-vary, measurements are correlated, and expression levels within a pathway tend to move together. When the baseline covariance is not the identity, the eigenvalues of the heterozygous covariance matrix reflect both these pre-existing correlations and any POE signal. The leading eigenvalue may be large simply because two phenotypes are strongly correlated, not because of a POE. This makes the simple-case estimator inapplicable directly (Figure 2, panel B).

**Fig. 2:**
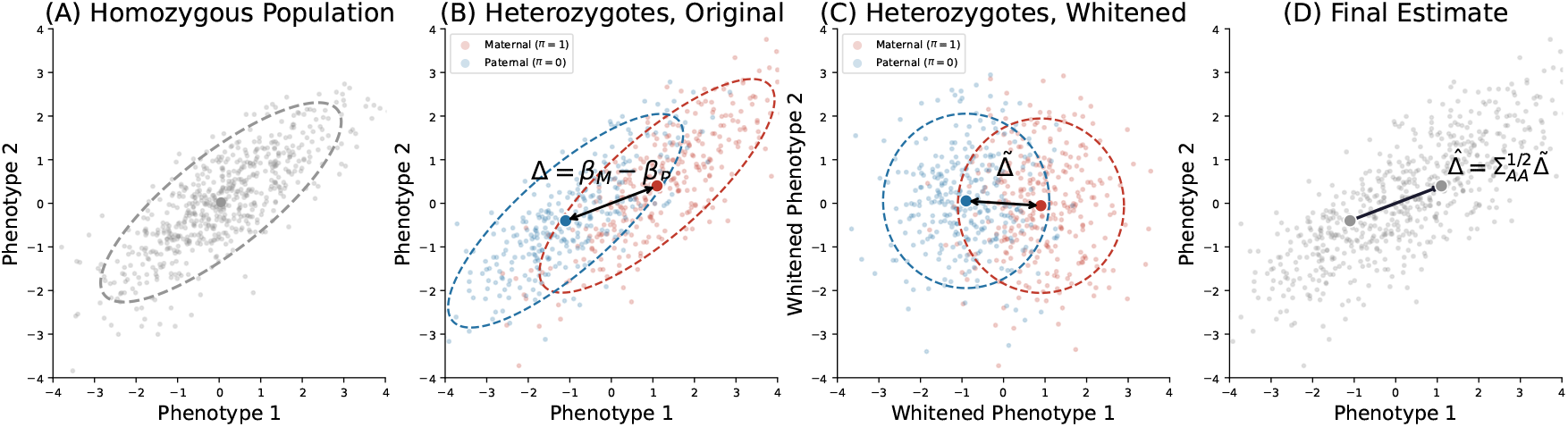
POE estimation with correlated phenotypes(general Σ_*AA*_, two synthetic phenotypes). **(A)** The homozygous (AA) population. These individuals share both alleles from the same reference type, so there is no parent-of-origin ambiguity. Their phenotype distribution provides an estimate of the baseline covariance 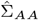. **(B)** The mean-centered heterozygous (AB) population. The true POE no longer aligns with the first principal component due to correlations between the phenotypes **(C)** The whitened heterozygous population, obtained by applying 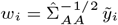 (Eq. 9). This transformation removes the baseline correlations, allowing the simple case to be used. **(D)** Finally the original estimate is recovered by undoing the transformation.

Our solution uses the homozygous population, which carries no POE signal (Figure 2, panel A), to measure and remove the baseline phenotype correlations before searching for the POE. The procedure has three steps:

1. **Estimate the baseline covariance from homozygotes**. We compute the empirical covariance of the mean-centered reference homozygotes,

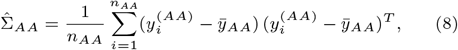

where 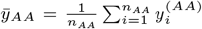. Because homozygotes carry no POE signal, 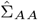 estimates the natural phenotype covariance uncontaminated by any parent-of-origin effect.
2. **Whiten the heterozygous data**. We transform the mean-centered heterozygous observations by

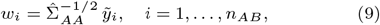

where 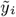 is the mean-centered heterozygous phenotype defined in Section 2.2.1. This “whitening” transformation rescales and decorrelates the phenotypes so that, in the transformed coordinate system, the within-group noise covariance is approximately the identity. The transformed data are now in the setting of Section 2.2.1, and the simple-case estimator applies.
3. **Estimate the POE in the whitened space, then transform back**. We apply equation (7) to the whitened data *{w*_*i*_*}* to obtain a POE estimate in the transformed space, 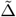. Because the whitening maps 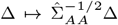, we recover the estimate in the original phenotype coordinates by reversing the transformation:

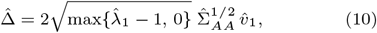

where 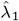 and 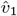 are the leading eigenvalue and eigenvector of the whitened heterozygous covariance. Multiplying by 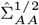 rotates and rescales the estimate back into the original phenotype space, where the entries of 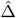 are directly procedure has three steps: interpretable as per-phenotype POE coefficients.

Figure 2 illustrates the full procedure on synthetic data with a POE in the first two correlated phenotypes: the homozygous population provides the baseline covariance (panel A); the raw heterozygous covariance conflates noise correlations with the POE (panel B); whitening removes the baseline structure so the excess variance from the POE becomes cleanly identifiable (panel C); and the final estimate, transformed back to the original scale, accurately recovers the true Δ (panel D).

### 2.3. Confidence Intervals via Bootstrap

We obtain confidence intervals on the POE estimates via a non-parametric bootstrap [Efron and Tibshirani, 1994]. Here we resample the data with replacement *B* times, and on each resampled dataset compute the POE estimate 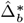. Because the sign of the leading eigenvector is not identifiable, we rectify each replicate so that its first entry is positive. The resulting collection 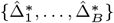 provides an empirical approximation to the sampling distribution of 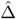. We therefore use the bias-corrected and accelerated (BCa) bootstrap [Efron, 1987], which adjusts the percentile endpoints to account for both bias and skewness, further detailed in the Appendix section A.4. For a fully distribution-free p-value a permutation test can be used as described in A.3 of the Appendix.

### 2.4. Minimum Detectable Effect Size and Filtering Criterion

With large samples a *p*-value can be small even when the effect is too weak to be reproducible, and model misspecification can produce false positives. We address both issues by requiring that any positive call correspond to a POE magnitude exceeding an information-theoretic detectability floor.

#### 2.4.1 Quartic scaling and Chernoff information

In a trio study, detecting a POE is a mean-shift test requiring *n* ∝ ∥Δ∥^−2^ observations. Without parental labels the signal enters only through the covariance perturbation ΔΔ^*T*^ */*4 (Eq. (5)), so resolving it against sampling fluctuations demands *n* ∝ ∥Δ∥^−4^. After whitening (Section 2.2.2), rotational symmetry (Proposition 1) reduces detection to a univariate test: *P*_0_ = *N* (0, 1) versus 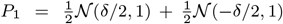, where 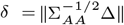 (Proposition 2). Proposition 5 (Appendix B) shows that the Chernoff information [Cover and Thomas, 2006] between these hypotheses is

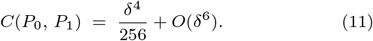

The quartic rather than quadratic leading order reflects the fact that *P*_0_ and *P*_1_ share the same mean; Theorem 4 (Appendix B) further establishes that this rate is minimax optimal.

#### 2.4.2. Minimum detectable effect size and filtering criterion

Defining *δ*^*\**^(*N, p, ϵ*) as the smallest POE for which the Chernoff-bounded Bayes error is at most *ϵ* [Cover and Thomas, 2006], we invert (11) to obtain

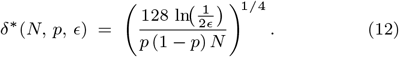

To integrate this threshold with the bootstrap uncertainty quantification of Section 2.3, we set *ϵ* = 0.001 and report a POE only if

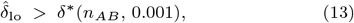

where 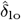 is the lower endpoint of the BCa confidence interval for each phenotype. This ensures that, even accounting for sampling variability, the estimated effect exceeds the information-theoretic detectability floor, excluding both practically insignificant effects and spurious signals from model misspecification.

## 3. Synthetic Examples

In this section, we analyze synthetic data focused on evaluating the Type 1 error rate, statistical power in terms of effect size, and robustness to model assumption violations. Note that the combination of Proposition 1 and Proposition 2 prove that the statistical properties of our estimator depend only on 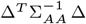. As a result, our simulations will assume Σ_*AA*_ = *I*.

### 3.1. Type 1 Error Rate

We first evaluate the performance of POISE under the null model. We conducted a simulation study with 8000 homozygotes and 2000 heterozygotes, *d* = 3 phenotypes, Σ_*AA*_ = *I*, and no marginal GWAS effect (*β* = *β* = 0). The data were generated from two noise distributions—a standard normal distribution and a *t*_9_ distribution—to evaluate sensitivity to heavy tails. We set the number of permutations to 10,000. Under the null hypothesis, a well-calibrated test should produce p-values that are approximately Uniform(0, 1). We generated 5,000 datasets with no true signal, applied the test to each replicate, and examined the resulting null p-values using a quantile–quantile (QQ) plot against the theoretical uniform distribution.

The results are shown in Figure 3. For both noise distributions, the points lie close to the *y* = *x* diagonal across the full range of quantiles, with no systematic upward deviation among small p-values and no broad curvature. We also compared the empirical p-value distributions to Uniform(0, 1) using a Kolmogorov–Smirnov test. The normally distributed data yielded a statistic of *D* = 0.029 with a p-value of 0.33, while the heavy-tailed data yielded a statistic of *D* = 0.039 corresponding to a p-value of 0.09, both strongly supporting a properly-calibrated test. We also emphasize that, in light of Proposition 1, this result extends to all non-singular Σ_*AA*_.

**Fig. 3:**
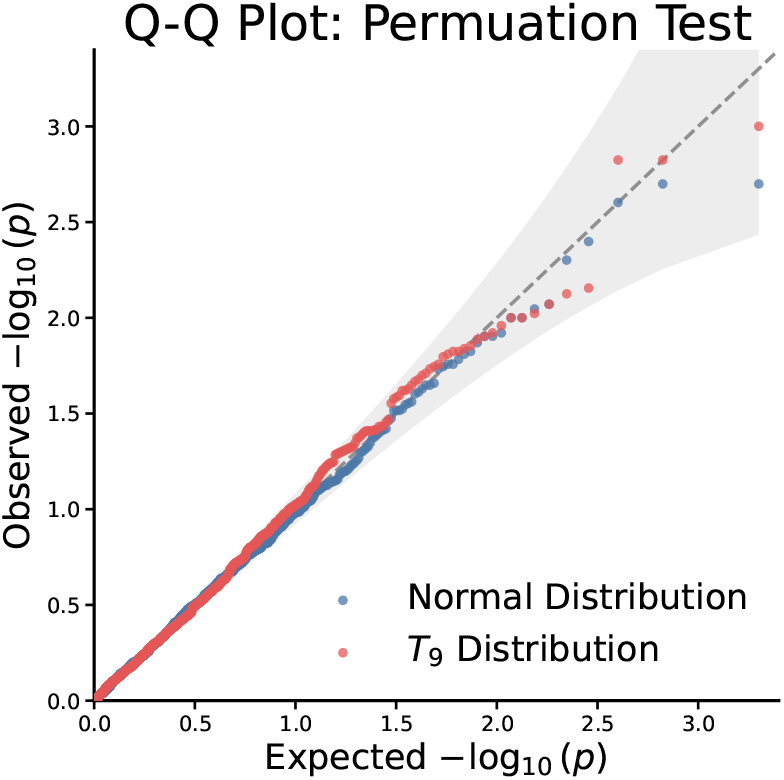
POISE maintains proper calibration under both Gaussian and heavy-tailed noise. QQ plot of empirical null p-values against the Uniform(0, 1) distribution for *d* = 3 phenotypes, 8,000 homozygotes, and 2,000 heterozygotes, based on 5,000 null replicates each.

**Fig. 4:**
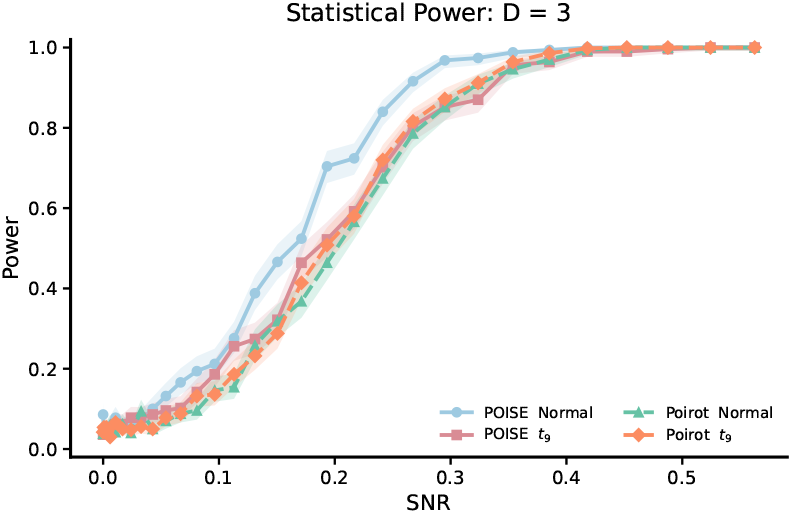
Statistical power of POISE and POIROT for detecting parent-of-origin effects under normal and heavy-tailed error distributions. Rejection rates at significance level *α* = 0.05 are plotted against signal-to-noise ratio (SNR = *κ*^2^) for a *p* = 3 dimensional phenotype with *N* = 2,000 subjects and minor allele frequency 0.25. Solid lines denote POISE; dashed lines denote POIROT. Shaded bands indicate 95% Wilson confidence intervals over 500 simulation replicates. Under Gaussian errors, POISE achieves substantially higher power than POIROT at moderate SNR, while under *t*_9_ errors the two methods perform comparably.

### 3.2. Power

To compare the statistical power of POISE and POIROT for detecting parent-of-origin effects in multivariate phenotypes, we conducted a simulation study under a mixture model for heterozygotes. We generated genotypes *g* ~ Binomial(2, 0.25) for *N* = 2,000 subjects with a *p* = 3 dimensional phenotype. Homozygous individuals (*g* = 0 or *g* = 2) were assigned phenotypes drawn from *N* (**0**, *I*_3_), while heterozygous individuals (*g* = 1) were drawn from a symmetric two-component mixture 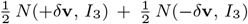 with **v** = (1, 0, 0)^*T*^, where *δ* controls the signal strength. We varied *δ* from 0 to 0.75 across 30 equally spaced values, corresponding to a signal-to-noise ratio SNR = *κ*^2^ ranging from 0 to 0.5625. To assess robustness to distributional misspecification, we repeated the simulation replacing normal errors with *t*_9_ errors rescaled to unit variance. For each setting, 500 independent replicates were performed, and rejection rates were computed at significance level *α* = 0.05, with POISE *p*-values obtained via 2,000 permutations.

Under normally distributed errors, POISE exhibited uniformly higher power than POIROT across the range of signal strengths examined. The advantage was most pronounced at moderate SNR: at *κ* = 0.44 (SNR = 0.19), POISE achieved a rejection rate of 70.4% compared with 46.4% for POIROT, a difference of 24 percentage points. POISE attained 80% power at SNR ≈ 0.24, whereas POIROT required SNR ≈ 0.29 to reach the same level; both methods converged to unit power beyond SNR ≈ 0.45. Under *t*_9_-distributed errors, the two methods exhibited comparable power throughout, with a mean absolute difference in rejection rate of only 0.02 across all SNR values and both reaching 80% power at SNR ≈ 0.27. As expected, heavy-tailed noise reduced power for both methods relative to the normal setting—for example, at *κ* = 0.49 (SNR = 0.24), power decreased from 84.0% to 70.2% for POISE and from 67.4% to 72.0% for POIROT—confirming that the Gaussian optimality of POISE does not come at the cost of robustness under distributional misspecification.

### 3.3. Robustness to vQTLs

There is one other aspect of POEs we wish to emphasize. We can see from (5) that the rank-one update to the covariance from a POE is positive-semidefinite. In other words, no additive POE can shrink the covariance of the heterozygotes compared to the homozygotes. The two-sample test underlying POIROT [Head et al., 2023] ignores this specific structure. As a result, it flags cases where 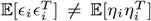, that is, when the within-group covariance structure differs between genotype groups, which can occur due to vQTLs [Dumitrascu et al., 2019]. This can be seen in cases where these tests flag variants where the variance is shrunk relative to the homozygous population.

In our third synthetic experiment, we contrast POIROT and POISE in the presence of such vQTLs. We violate the shared-covariance assumption 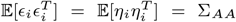 by setting 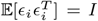 and 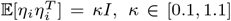, uniformly scaling the heterozygote covariance relative to the homozygous baseline. We set *N* = 10000, *p* = 3, and MAF of 0.25, repeating each simulation 100 times to obtain confidence intervals on the rejection rate. Critically, when *κ <* 1, no additive POE can account for the data. Because additive POE contributes a positive-semidefinite rank-one term to the heterozygote covariance, it can only increase variance relative to the homozygous baseline. A uniform shrinkage with *κ <* 1 is therefore incompatible with any additive POE.

Figure 5 plots the results. POIROT rejects the null consistently whenever *κ* ∉ [0.95, 1.05], which is expected from the three-sample covariance test it employs: at *N* = 10000, that test achieves power near 1 against any detectable covariance difference, regardless of whether the deviation is compatible with an additive POE. POISE, in contrast, has zero false positives for all *κ <* 1. This immunity follows directly from the eigenvalue truncation in the estimator: asymptotically, *λ*_1_ = *κ <* 1 when *κ <* 1, so 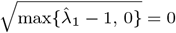, which forces 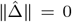 and a permutation *p*-value of 1. We also note that this robustness is one-sided. The eigenvalue truncation 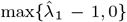 prevents false positives when the heterozygote covariance is *deflated* (*κ <* 1), since the leading eigenvalue of the whitened covariance falls below 1 and the estimator returns zero. When *κ >* 1 the leading eigenvalue exceeds 1 regardless of whether the excess variance has the rank-one mixture structure of an additive POE or is a full-rank vQTL perturbation, and POISE yields a nonzero estimate.

**Fig. 5:**
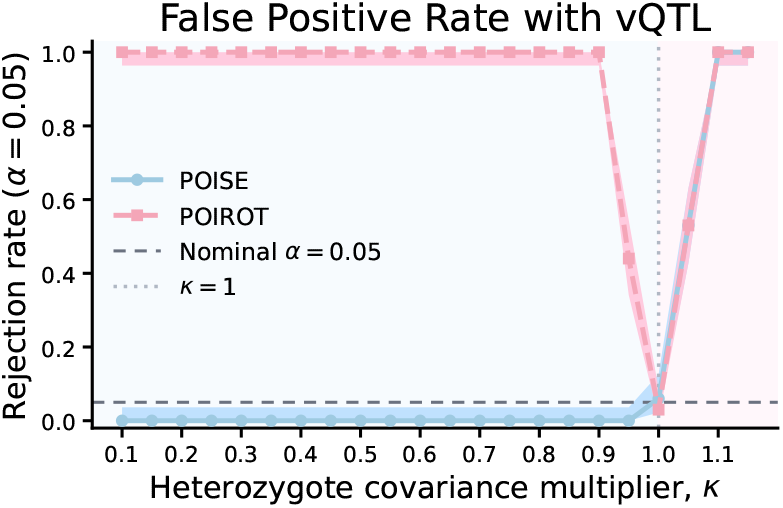
POISE maintains zero false positives under a variance-shift null; POIROT does not. False-positive rates for POISE and POIROT as the heterozygote covariance multiplier *κ* varies over [0.1, 1.1] under a pure variance-shift null (no true parent-of-origin effect). When *κ <* 1, a uniform covariance shrinkage in the heterozygote group is incompatible with any additive parent-of-origin effect, yet POIROT rejects broadly; POISE produces zero false positives throughout, owing to the eigenvalue truncation 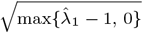. Points and shaded bands show estimated rejection rates and 95% Clopper-Pearson confidence intervals across 100 simulation replicates.

**Fig. 6:**
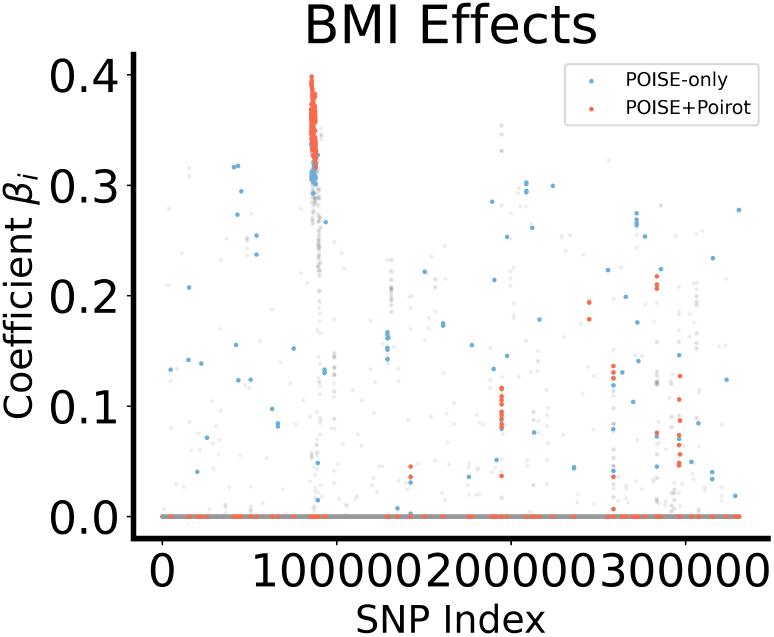
POE estimates from the UK Biobank for the BMI phenotype. This plot is similar to a Manhattan plot, except it plots the rectified coefficient rather than a log p-value. Coefficients not significant according to both POIROT and POISE are set to 0. Statistically significant coefficients that fail the practical significance test are plotted in grey. Coefficients that are practically significant and detected by both POIROT and POISE are shown in red. Blue points are SNPs detected exclusively by POISE, illustrating the broader set of signals recovered by our method.

## 4. UK Biobank Results

### 4.1. Procedure

We analyzed genetic and phenotypic data from 602,000 participants in the UK Biobank [Sudlow et al., 2015], a large-scale biomedical cohort designed to investigate genetic and environmental determinants of health in individuals aged 40–69 from across the UK. The participants provided a range of biological samples, such as blood, urine, and saliva, as well as detailed health and lifestyle information through questionnaires and physical measurements. We focused on body mass index (BMI), LDL cholesterol, and HDL cholesterol. These traits were chosen as they are complex with prior links to POEs, and to facilitate comparison with POIROT [Head et al., 2023]. We estimated the POEs with these three features for every variant with MAF greater than 0.05, after adjusting for genotype array, ancestry PCs, sex, age, and smoking status. We prescreened variants using Bonferroni-corrected p-values derived from POIROT [Head et al., 2023] and the two-sample test from [Zheng et al., 2015]. We found that the latter method was not robust to deviations from normality, as a result we used an inverse normal transformation on all phenotypes.

At a Bonferroni threshold of 1.5 *×* 10^−7^, POIROT identified 338 variants, of which 186 met the effect size criterion (Section 2.4). POISE identified 893 variants, of which 320 passed the same criterion. Critically, all 186 POIROT-significant variants were contained within the POISE set. In other words, POISE is a strict superset of POIROT with respect to POEs with large effect estimates. POISE additionally identified 134 variants that POIROT did not detect. To validate that POISE recovers genuine signals, we first examined the 186 variants detected by both methods. This concordant set serves as a sensitivity benchmark: because POIROT is a conservative, previously validated test, the complete overlap confirms that POISE does not miss true positives.

The concordant set was dominated by a signal on chromosome 6: 152 of 186 variants (81.7%) mapped to the extended MHC class I region at 6p22.1–p21.32, spanning approximately 5 Mb (hg19 positions 25.9–30.9 Mb). The remaining 34 variants were distributed across six other chromosomes, with chromosome 11 (*n* = 13), chromosome 19 (*n* = 8), chromosome 16 (*n* = 5), chromosome 2 (*n* = 4), chromosome 15 (*n* = 2), and chromosome 8 (*n* = 2). The concordant set was characterized by relatively low population allele frequencies (mean gnomAD genomes global AF = 0.098; NFE AF = 0.109), with particularly pronounced rarity in East Asian populations (mean EAS AF = 0.033), and carried larger absolute effect sizes on BMI than the POISE-exclusive variants (mean |*β*| = 0.312 vs. 0.208).

Outside of chromosome 6, the concordant variants encompassed several canonical lipid-metabolism GWAS loci with established relevance to cardiometabolic traits. These included four variants at the *APOB* locus on chromosome 2—including the nonsynonymous variant rs1367117 (p.Thr98Ile; CADD = 22.0), a well-established LDL cholesterol locus (GWAS *p* = 4.48 *×* 10^−114^) for which [Hochner et al., 2015] reported a maternal-specific effect on BMI (*β* = 0.8 kg/m^2^; *p* = 3.1 *×* 10^−5^) and waist circumference (*β* = 2.7 cm; *p* = 2.1 *×* 10^−7^). On chromosome 11q23.3, 13 variants mapped to the *APOA5* /*APOC3* /*BUD13* /*ZPR1* apolipoprotein gene cluster, including the functional missense variant rs3135506 (*APOA5* p.Ser19Trp; CADD = 23.7), which reduces apolipoprotein A-V secretion by approximately 50% and is a major triglyceride and HDL-cholesterol locus. An additional eight variants at the *APOE* /*TOMM40* locus on chromosome 19, five variants at the *CETP* locus on chromosome 16, and two variants near *LIPC* on chromosome 15 completed a comprehensive representation of the established lipid-metabolism genetic architecture.

Overall, the concordant set was markedly enriched for coding variants relative to the POISE-exclusive group, with 20 exonic variants (10.8% of 186) compared to two in the exclusive set (1.5% of 133). Eight variants exceeded the CADD ≥ 15 threshold for potential deleteriousness, and six exceeded CADD ≥ 20. Six variants carried ClinVar annotations: rs1367117 (*APOB*; linked to familial hypercholesterolemia), rs3135506 (*APOA5*; hypertriglyceridemia and cardiovascular phenotype), rs769449 (*APOE*), rs1800588 (*LIPC*; HDL-cholesterol QTL12, classified as “association”), rs1864163 (*CETP*), and rs2072803 (*BTN2A2*).

Beyond the concordant set, POISE identified 134 additional SNPs that POIROT did not detect. These variants constitute the primary evidence that POISE offers superior detection power. The POISE-exclusive set exhibited a substantially broader genomic distribution than the concordant variants, spanning 19 chromosomes, with chromosome 6 still the most represented (*n* = 51; 38.3%) but with appreciable contributions from chromosomes 8 (*n* = 12), 17 (*n* = 9), 1 (*n* = 8), 2 (*n* = 8), 11 (*n* = 8), 4 (*n* = 7), 3 (*n* = 5), and 16 (*n* = 5), among others. The allele frequency profile was significantly higher than the concordant set, with a mean gnomAD genomes global AF of 0.218 (median 0.196) and mean NFE AF of 0.232, indicating that POISE captures common-variant signals that lack the heterozygote-variance signature detected by POIROT. The POISE-exclusive variants carried smaller absolute effect sizes (mean |*β*| = 0.208) and more moderate *p*-values (median *p* = 1.4 *×* 10^−4^ vs. 1.0 *×* 10^−5^ in the concordant set), consistent with the expectation that POISE’s enhanced sensitivity extends detection to more subtle effects.

While not a comprehensive analysis of biological relevance, many of the novel POISE-exclusive variants mapped to canonical obesity and metabolic GWAS genes. These included rs12243326 in *TCF7L2* (chr10), the most strongly replicated type 2 diabetes susceptibility gene, which was also reported as a genome-wide significant BMI locus (*p* = 1.11*×*10^−11^) in the GIANT consortium meta-analysis [**?**]. The POISE-exclusive set also contained variants at well-established LDL cholesterol loci. Two variants in *ABCG8* on chromosome 2 implicate the ABCG5/ABCG8 sterol transporter complex, which limits intestinal cholesterol absorption and promotes biliary secretion. The ABCG5/ABCG8 locus is among the most significant in the Global Lipids Genetics Consortium (GLGC) meta-analysis, with LDL-C association *p* = 2 *×* 10^−47^ [**?**]. POISE also identified rs10846744 in *SCARB1* on chromosome 12, encoding the HDL scavenger receptor SR-BI, which mediates selective uptake of HDL cholesteryl ester and is a genome-wide significant HDL-C locus (*p* = 3 *×* 10^−14^ in GLGC) [**?**]. Additionally, rs708272 in *CETP*, the classic TaqIB polymorphism associated with a per-allele HDL-C increase of approximately 3.1 mg/dL [**?**], was detected exclusively by POISE (classified as benign for coronary artery disorder in ClinVar). Together with the concordant-set signals at *CETP, LIPC*, and the 11q23.3 apolipoprotein cluster, these POISE-exclusive variants demonstrate that POISE captures a more comprehensive representation of the HDL-C genetic architecture than POIROT alone.

## 5. Conclusion

We present POISE, a spectral framework for detecting parent-of-origin effects in GWAS data from unrelated individuals. By modeling the heterozygous population as an unobserved mixture and recovering the POE vector from the leading eigenvector of the whitened covariance, POISE provides per-trait effect-size estimates with bootstrap confidence intervals, capabilities not available from existing GWAS-based POE methods. The information-theoretic minimum detectable effect size gives researchers a closed-form criterion for assessing whether a given sample is large enough to resolve a POE of a given magnitude, and the permutation test provides exact Type I error control without distributional assumptions.

Our approach is complementary to phasing-based methods that directly assign parental origin [Hofmeister et al., 2022, 2025]. Those methods yield the most direct evidence for POE, including the direction of the parental effect, and are not susceptible to confounding by non-POE sources of variance heterogeneity. However, they require identifiable surrogate relatives and are limited to the fraction of a cohort for which parental-origin inference is feasible. POISE operates on the full cohort without phasing, extending POE screening to settings where relatedness structure is sparse or absent, and exploiting the larger effective sample size to detect effects that fall below the power threshold of smaller phasing-eligible subsets.

Several limitations of the present work should be noted. First, like all variance-based POE methods, POISE cannot distinguish a true parent-of-origin effect from an inflation-type variance QTL arising from gene–environment interaction, epistasis, or other mechanisms. Second, the direction of the parental effect (maternal vs. paternal) is fundamentally unidentifiable from unlabeled heterozygotes, so POISE findings at novel loci require follow-up with phasing-based or family-based methods to determine the biological direction of the effect.

## 8. Data Availability

The code for this method in Python is publicly available at https://github.com/bystrogenomics/POISE.

## A Properties of the Spectral POE Estimator

### A.1 Mathematical Derivations

In this section we prove several claims about our spectral POE estimator: that it is rotationally equivariant (with an invariant *ℓ*_2_ test statistic), that its statistical properties depend solely on the Mahalanobis SNR, and that the Mahalanobis SNR dominates the directional SNR.

#### Proposition 1

(Rotational equivariance) *Let Q* ∈ *O*(*d*) *be an arbitrary orthogonal matrix (that is Q*^*T*^ *Q* = *I*) *and define rotated data z*_*i*_ = *Qy*_*i*_ *for every individual (both homozygotes and heterozygotes). Denote by* 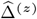 *and* 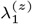 *the POE estimate and largest eigenvalue of the data {z*_*i*_*}, and by* 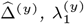 *the values obtained from the original data. Then:*

1. 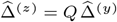 (*equivariance of the estimate*);
2. 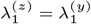^)^ (*invariance of the test statistic*);
3. *the permutation p-value based on the ℓ*_2_ *norm* 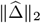 *is invariant*.

*Proof* First note that Cov(*AX*) = *A*Cov(*X*)*A*^*T*^ for any variable *X* and matrix *A*. As a result, the covariance of the rotated data is

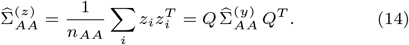

Next, for any positive-definite *A* and orthogonal *Q*, the unique positive-definite square root satisfies (*QAQ*^*T*^)^1*/*2^ = *Q A*^1*/*2^ *Q*^*T*^, since

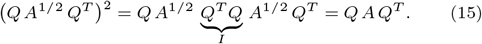

Applying this to (14):

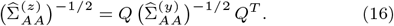

This is the new whitening transformation on the rotated homozygous population. When this is applied to the rotated heterozygotes we obtain

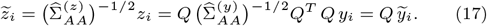

Using the same properties as before, the new covariance matrix of the whitened heterozygotes is

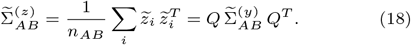

We now derive the relationship between the eigenvalues and eigenvectors of 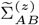 and 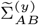. Suppose 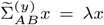 for some vector *x*. Multiplying on the left by *Q* and setting *a* = *Qx* gives

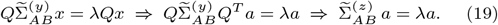

Hence all eigenvalues of 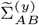 and 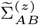 are shared, and in particular 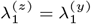. Moreover, if 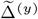 is the leading eigenvector of 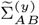, then 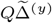 is the leading eigenvector of 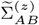, so

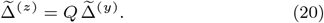

Substituting these quantities into the estimator formula (10) yields

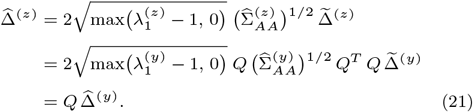

This proves 1 and 2.

Finally, since *Q* is orthogonal, 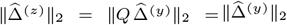. The same equivariance holds for every permuted replicate: 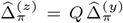 for any permutation *π*, so 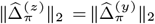. Hence the rank of the observed statistic within the permutation null distribution is unchanged, proving 3. □

Note that the *ℓ*_1_ and *ℓ*_*∞*_ norms are *not* rotationally invariant: ∥*Qx*∥_1_ ≠ ∥*x*∥_1_ and ∥*Qx*∥_*∞*_ ≠ ∥*x*∥_*∞*_ in general. Our next step is proving that the statistical properties of our estimator and statistical test depend only on a single quantity, the signal-to-noise ratio (SNR).

#### Proposition 2

(Reduction to Mahalanobis SNR) *At the population level* 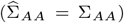, *all statistical properties of the estimator, including the type I error, power, bootstrap coverage, bias, and variance of* ∥Δ∥_2_, *depend on* (Σ_*AA*_, Δ) *only through the scalar*

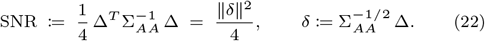

*Proof* We first whiten the heterozygous population using the homozygous covariance matrix. Writing *µ* = *δ/*2, a draw from the mixture 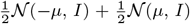 satisfies 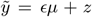, where *ϵ* ~ Uniform(*{*−1, +1*}*) is independent of *z* ~ *N* (0, *I*). Using 𝔼[*ϵ*^2^] = 1 and 𝔼[*ϵ*] = 0, the second moment is

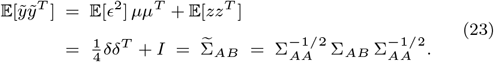

This is a rank-1 spiked identity model with eigenvalues *λ*_1_ = 1 + ∥*δ*∥^2^*/*4 (eigenvector *δ/*∥*δ*∥) and *λ*_2_ = *· · ·* = *λ*_*d*_ = 1, as mentioned in Section 2.2.1.

Since all statistics of interest are functions only of the sample covariance

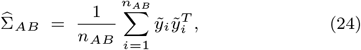

it suffices to show that the distribution of 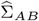 depends on (Σ_*AA*_, Δ) only through ∥*δ*∥^2^ = 4 SNR. The finite-sample distribution of 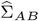 is determined by the moments of 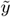; we show each moment depends on *δ* only through ∥*δ*∥^2^ by establishing that the full distribution of 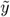 is invariant, up to rotation, under changes to *δ* that preserve ∥*δ*∥^2^.

Using the decomposition 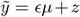 and expanding, the fourth-moment tensor of 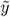 is

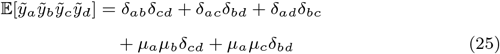

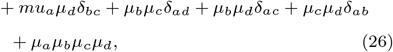

where we used 𝔼[*ϵ*^*k*^] = 1 for *k* even and 0 for *k* odd, together with the Gaussian moment identity 𝔼[*z*_*a*_*z*_*b*_*z*_*c*_*z*_*d*_] = *δ*_*ab*_*δ*_*cd*_ + *δ*_*ac*_*δ*_*bd*_ + *δ*_*ad*_*δ*_*bc*_. Every term in (25) is a product of factors drawn from *{δ*_*ab*_, *µ*_*a*_*}* = *{δ*_*ab*_, *δ*_*a*_*/*2*}*, so the entire tensor is a function of *µ* = *δ/*2 through the rank-1 matrix *µµ*^*T*^ = *δδ*^*T*^ */*4 and the identity. The same holds for all higher moments by an identical parity argument: odd powers of *ϵ* vanish, so every moment of 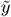 is a polynomial in *µµ*^*T*^ and *I*.

Now let *Q* ∈ *O*(*d*) be any orthogonal matrix and consider replacing *δ* → *Qδ*, equivalently *µ* → *Qµ*. Under this substitution, 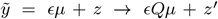 where *z*^*′*^ = *Qz* ~ *N* (0, *I*), so the transformed variable equals 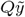 in distribution. Consequently, the distribution of 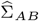 under spike direction *Qδ* equals the distribution of 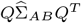 under spike direction *δ*. Since ∥*Qδ*∥^2^ = ∥*δ*∥^2^ for any orthogonal *Q*, the orbit of *δ* under *O*(*d*) is the sphere ∥*δ*∥ = const, and the distribution of 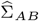 is constant on this orbit. Therefore every statistic that is invariant under 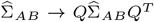, including the eigenvalues of 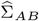, the norm 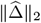, and the permutation *p*-value under the *ℓ*_2_ norm, depends on *δ* only through ∥*δ*∥^2^ = 4 SNR. □

#### Proposition 3

(Comparison with directional SNR) *Let u* = Δ*/*∥Δ∥ *and define the* directional signal-to-noise ratio

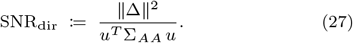

*Then*

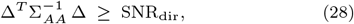

*with equality if and only if u is an eigenvector of* Σ_*AA*_.

*Proof* Write 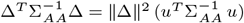, so the inequality (28) is equivalent to

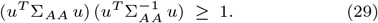

Let Σ_*AA*_ = *U* Λ*U* ^*T*^ be an eigendecomposition with Λ = diag(*ℓ*_1_, …, *ℓ*_*d*_), *ℓ*_*j*_ *>* 0, and set *w* = *U* ^*T*^ *u* (so ∥*w*∥ = 1). Then

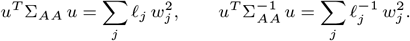

By the Cauchy–Schwarz inequality applied to the vectors 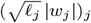 and 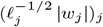:

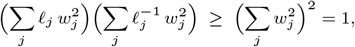

with equality if and only if 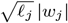 is proportional to 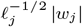 for all *j*, i.e., 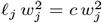 for some constant *c* whenever *w*_*j*_≠ 0.

This holds precisely when all nonzero *w*_*j*_ correspond to a single eigenvalue, i.e., *u* lies in an eigenspace of Σ_*AA*_. □

Note that the Mahalanobis SNR is at least as large as the directional SNR because the whitening transformation uses the full covariance structure, not merely the variance along Δ. When noise is large along Δ but small in orthogonal directions, whitening compresses the noisy direction and expands the quiet ones, effectively redistributing the signal favorably.

### A 2. Simulation Validation

In each demo the heterozygous observations are drawn from the symmetric mixture

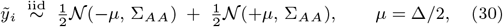

implemented as 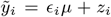 where *ϵ*_*i*_ ~ Uniform(*{*−1, +1*}*) is independent of *z*_*i*_ ~ *N* (0, Σ_*AA*_). The Cholesky factor of Σ_*AA*_ is used to generate *z*_*i*_.

#### A.2.1 Reduction to Mahalanobis SNR (Proposition 2)

We verify that two settings with the same Mahalanobis SNR but different (Σ_*AA*_, Δ) yield the same distribution of 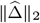. We take *d* = 8, *n*_*AB*_ = 400, and *n*_reps_ = 2,000 Monte Carlo repetitions.

*Setting A* uses Σ_*A*_ = *I*_*d*_ and Δ_*A*_ = 2 SNR *e*_1_, giving SNR = ∥Δ_*A*_∥^2^*/*4.

*Setting B* uses an anisotropic Σ_*B*_ = *Q* diag(*ℓ*_1_, …, *ℓ*_*d*_) *Q*^*T*^ with *Q* a random orthogonal matrix and eigenvalues *ℓ*_*j*_ equally spaced in [0.5, 2.0]. The signal is constructed as 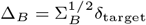 where 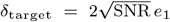, so that 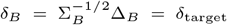 and ∥*δ*_*B*_∥^2^ = ∥*δ*_*A*_∥^2^, matching the SNR exactly. The target SNR is set to 1.5 in all runs. If Proposition 2 holds, the empirical distributions of 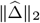 under settings A and B should be indistinguishable in Figure 7.

**Fig. 7:**
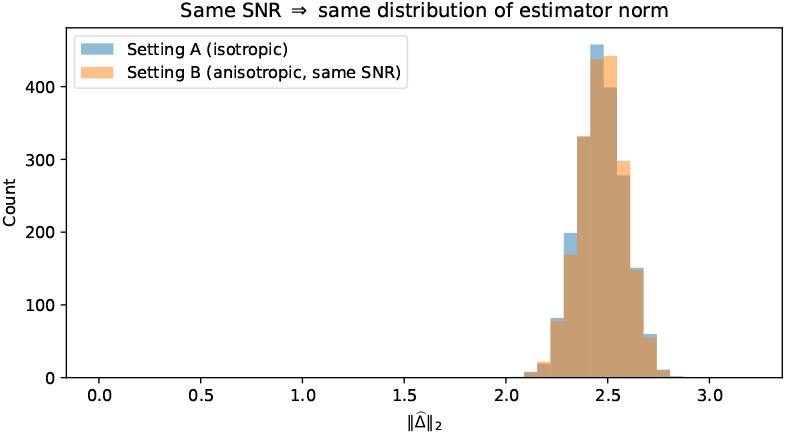
Equal Mahalanobis SNR implies equal estimator distribution. Empirical distributions of 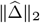 over 2,000 Monte Carlo replicates for two settings with identical SNR = 1.5 but different (Σ_*AA*_, Δ): Setting A (isotropic, Σ_*A*_ = *I*) and Setting B (anisotropic, Σ_*B*_≠ *σ*^2^*I*, constructed to match the Mahalanobis SNR exactly). The near-perfect overlap of the two histograms confirms that all distributional properties of the estimator depend on (Σ_*AA*_, Δ) only through 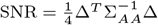 (Proposition 2).

#### A.2.2. Demo 2: Rotational invariance (Proposition 2)

We verify that rotating *δ* by an arbitrary orthogonal matrix *Q* leaves the distribution of 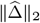 unchanged. We take Σ_*AA*_ = *I*_*d*_ (so *δ* = Δ), *d* = 6, *n*_*AB*_ = 300, *n*_reps_ = 2,000, and *δ*_base_ = 2*e*_1_ (SNR = 1). Three spike directions are compared: the original *δ*_base_ and two random rotations *Q*_1_*δ*_base_, *Q*_2_*δ*_base_, where *Q*_1_, *Q*_2_ ∈ *O*(*d*) are drawn independently. Since ∥*Q*_*k*_*δ*_base_∥^2^ = ∥*δ*_base_∥^2^ for all *k*, Proposition 2 predicts identical distributions across all three directions, as visualized in Figure 8.

**Fig. 8:**
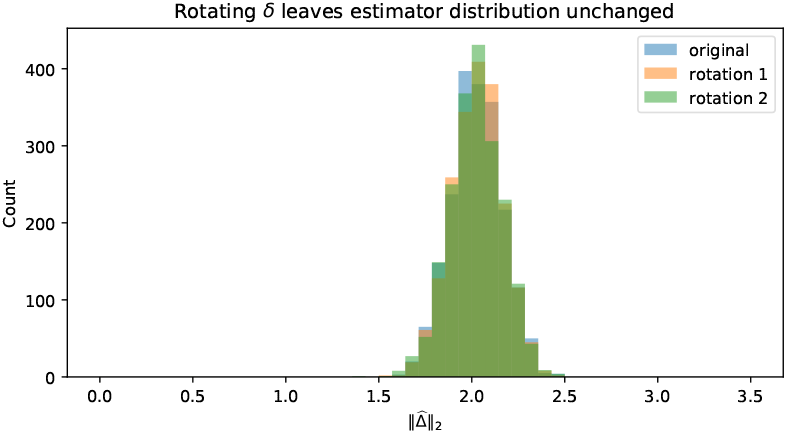
Rotating *δ* leaves the estimator distribution unchanged. Empirical distributions of 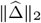 over 2,000 Monte Carlo replicates for three spike directions: the canonical direction *δ*_base_ = 2*e*_1_ and two random orthogonal rotations *Q*_1_*δ*_base_, *Q*_2_*δ*_base_. All three directions share the same norm ∥*δ*∥ = 2 (SNR = 1). The three histograms are indistinguishable, confirming the rotational invariance of the estimator (Proposition 2): any rotationally invariant functional of 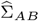 depends on *δ* only through ∥*δ*∥^2^.

### A.3. P-value Estimation via a Permutation Test

While the POE estimates and confidence intervals from Sections 2.2 and 2.3 are informative, evaluating their statistical significance is critical, especially given that spectral inflation, the tendency of leading sample eigenvalues to exceed their population counterparts at finite sample sizes [Ledoit and Wolf, 2004], introduces an upward bias in 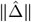. Unfortunately, the estimator’s reliance on matrix inversion, spectral decomposition, and shrinkage complicates the derivation of an analytic sampling distribution.

However, the structure of our estimator is well suited to a permutation test. Under the null hypothesis *H*_0_ : Δ = 0, the heterozygous phenotype model reduces to 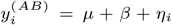 with Cov(*η*_*i*_) = Σ_*AA*_, a single population with no inheritance-dependent substructure. After mean-centering each genotype group, both the homozygous and heterozygous observations are draws from the same zero-mean distribution with covariance Σ_*AA*_. The genotype group labels *AA* and *AB* are therefore exchangeable under *H*_0_: any reassignment of the *n*_*AA*_ + *n*_*AB*_ mean-centered observations into two groups of sizes *n*_*AA*_ and *n*_*AB*_ is equally likely. This exchangeability makes the permutation test exact: the resulting *p*-value has the correct Type I error rate under the null by construction, without requiring asymptotic or parametric assumptions.

To perform the test, we randomly reassign the genotype group labels across the *n*_*AA*_ + *n*_*AB*_ mean-centered individuals, run the full estimation pipeline (whitening, spectral decomposition, back-transformation) on the permuted data, and compute a test statistic. We use the norm of the POE estimate,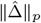, as the test statistic, where *p* = 1, 2, *∞* correspond to the *ℓ*_1_, *ℓ*_2_, and maxnorms, respectively. Repeating this for *R* random permutations produces a null distribution 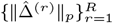, and the *p*-value is the fraction of permutations for which the permuted test statistic meets or exceeds the observed value. We recommend *ℓ*_2_, as the resulting *p*-value is invariant under arbitrary correlation structures in Σ_*AA*_; we prove this mathematically in the Appendix Section A.1 and demonstrate it empirically in Section A.2.

An important property of the permutation test is that the *p*-value remains valid even when the point estimate 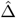 is biased symmetric (*z*_0_ = *a* = 0), this reduces to the ordinary percentile in magnitude. Crucially, this bias affects the observed test statistic and every permuted test statistic in exactly the same way, because the identical estimation pipeline is applied to each permutation. Consequently, the relative ranking of the observed statistic within the permutation null distribution is unaffected by spectral inflation, and the *p*-value is determined solely by this ranking, which we demonstrate in Section 3.1. In particular, when Δ = 0 the observed and permuted datasets are drawn from the same distribution, so the observed statistic has no systematic tendency to rank high regardless of any shared upward bias. A truly null variant will therefore yield a non-significant *p*-value even if 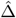 substantially overestimates ∥Δ∥ in absolute terms.

Although our estimation algorithm is fast, typically under one second per variant, even a modest number of permutations creates a bottleneck at GWAS scale, where hundreds of thousands of variants must be tested. We propose two strategies to address this. The first is to use an analytical hypothesis test, such as those of Zheng et al. [2015] or Head et al. [2023], as a preliminary screening step to flag candidate variants for the more computationally intensive permutation procedure. The second strategy exploits a key feature of our framework: although genotype groupings vary across loci, the *d*-dimensional phenotype matrix remains fixed. This enables the reuse of permutation indices across variants with similar minor allele frequencies (MAFs). In large-scale studies, a one-time permutation cost can thus be amortized across thousands of tests. However, the null distribution of 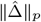 is sensitive to the relative sizes of the homozygous and heterozygous groups, so sufficient permutation density is still required to accurately cover the range of *n*_*AA*_*/n*_*AB*_ ratios observed across loci.

### A.4 BCA-Based Confidence Intervals

Working phenotype-by-phenotype, let 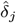 denote the point estimate of *δ*_*j*_ = Δ_*j*_. The BCa method requires two correction factors. The *bias correction z*_0_ measures how the bootstrap distribution is centered relative to the point estimate:

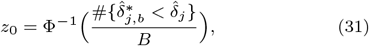

where Φ is the standard normal CDF. The *acceleration a* captures the rate at which the standard error of 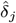 changes with the true parameter value, estimated via the jackknife [Miller, 1974]:

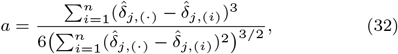

where 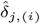 is the estimate obtained by omitting the *i*-th observation and 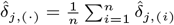 is the jackknife mean. The BCa interval then replaces the nominal percentiles *α/*2 and 1−*α/*2 with the adjusted endpoints

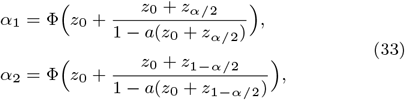

where *z*_*q*_ = Φ^−1^(*q*), and reads off the confidence interval as 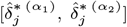. When the bootstrap distribution is unbiased and symmetric (z_0_ = a = 0), this reduces to the ordinary percentile interval.

The jackknife required by the acceleration factor is the dominant computational cost: for biobank-scale data it requires re-estimating 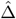 once per heterozygous individual. To make this tractable, we exploit the structure of our estimator, which separates the estimation of the whitening transformation (from homozygotes) from the estimation of the POE signal (from heterozygotes). Because the whitening step depends only on the homozygous population and is comparatively stable, we fix 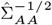 at its point estimate and restrict both the bootstrap and jackknife to resampling the whitened heterozygous observations only. This reduces the jackknife from *n*_*AA*_ + *n*_*AB*_ leave-one-out repetitions to *n*_*AB*_ alone, typically yielding one to two orders of magnitude improvement in computation time without appreciable loss in coverage accuracy.

## B. Minimax Lower Bounds: POISE Optimality

### B.1. Mathematical Derivations

We now establish that the estimation error of any POE estimator must be at least of order 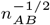 at small effect sizes. Unlike most standard parametric estimation, the POE signal Δ enters the distribution of heterozygous observations only through the fourth cumulant: the *δ*^2^ perturbation to the density ratio *p*_Δ_*/p*_0_ is proportional to the second Hermite polynomial *H*_2_(*y*) = *y*^2^ − 1, which integrates to zero against *p*_0_ by orthogonality; consequently the Chernoff information between *p*_0_ and *p*_Δ_ scales as *δ*^4^ rather than *δ*^2^.

This quartic scaling shifts the minimax rate from the usual *n*^−1^ to *n*^−1*/*2^

Throughout, *p*_0_ = *N* (0, *I*) denotes the null (no-POE) density and

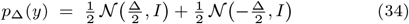

denotes the heterozygous mixture density under POE vector Δ ∈ R^*d*^. Write 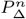 for the joint law of *n* = *n*_*AB*_ i.i.d. draws from *p*_Δ_.

The Chernoff information between two densities *p* and *q* is

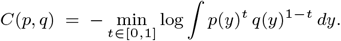

#### Theorem 4

(Minimax lower bound) *Fix d* ≥ 1. *There exists a universal constant c*_1_ *>* 0 *such that, for all n*_*AB*_ ≥ 1,

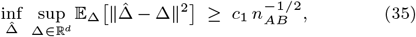

*where the infimum is over all estimators* 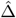 *based on n*_*AB*_ *i*.*i*.*d. observations from p*_Δ_.

The proof rests on two ingredients: a precise computation of the Chernoff information between *p*_0_ and *p*_Δ_ (Proposition 5), and Le Cam’s two-point method.

#### Proposition 5

(Chernoff information scales quartically) *For* Δ ∈ R^*d*^ *with* ∥Δ∥ = *δ*,

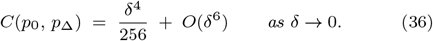

*In particular, C*(*p*_0_, *p*_Δ_) ≍ *δ*^4^ *for δ sufficiently small*.

*Proof* Due to the previously-established rotational symmetry, we can assume that all variation occurs in the first dimension via a change of coordinates. As such, we can take *d* = 1 and Δ = *δ >* 0. The Chernoff information is *C* = − min_*t*_ log *Z*(*t*) where

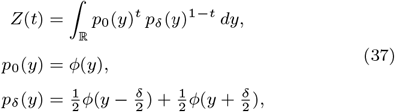

and *ϕ* is the standard Gaussian density. We first compute the Taylor expansion of *p*_*δ*_. Writing 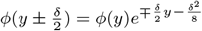,

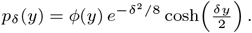

Expanding the exponential and cosh to sixth order in *δ*,

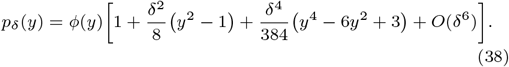

Crucially, the coefficient of *δ*^2^ is proportional to the second Hermite polynomial *H*_2_ (*y*) = *y*^2^ − 1, which is orthogonal to 1 under *ϕ*. Hence ∫ *p*_*δ*_(*y*) *dy* = 1+*O*(*δ*^4^), confirming that the leading deviation from *p*_0_ is of order *δ*^4^.

We then expand *Z*(*t*). Let *p*_*δ*_ = *p*_0_(1 + *f*_*δ*_) where *f*_*δ*_ = *O*(*δ*^2^) from (38). Then

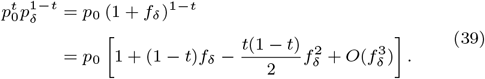

Integrating and using ∫ *p*_0_ *f* = *O*(*δ*^4^) (from the orthogonality noted above) and *p*_0_*f* ^2^ = *O*(*δ*^4^), we obtain

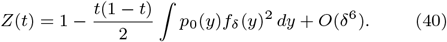

To evaluate the integral, the leading term of *f*_*δ*_ from (38) is 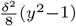, so

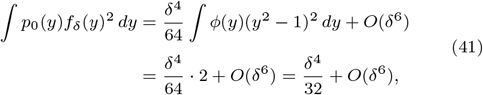

where we used ∫ *ϕ*(*y*)(*y*^2^ − 1)^2^ *dy* = y = 𝔼[(*Z*^2^ ™ 1)^2^] = 2 for *Z* ~ *N* (0, 1). Substituting back,

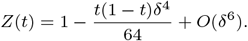

We then minimize over *t*. The function *t 1*→ *t*(1−*t*) is maximised at 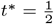 with value 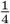, giving

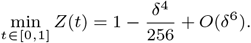

Hence

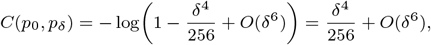

establishing (36). □

*Proof of Theorem 4* We apply Le Cam’s two-point method. Choose Δ_0_ = 0 and Δ_1_ = *δe*_1_ where *e*_1_ is the first standard basis vector and *δ >* 0 is to be optimised. For any estimator 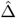,

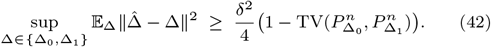

This is Le Cam’s inequality: the worst-case squared error is at least the squared separation times the probability that no test can distinguish the two hypotheses [LeCam, 1973, Yu, 1997].

It remains to lower-bound 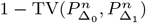. By the relation between total variation and Chernoff information,

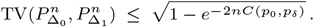

From Proposition 5, *C*(*p*_0_, *p*_*δ*_) ≤ *c*_0_*δ*^4^ for a constant *c*_0_ *>* 0 and all *δ* sufficiently small. Choose

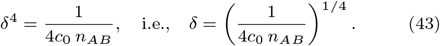

Then 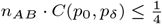, so

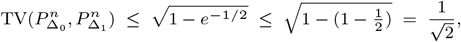

and hence 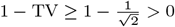.

Substituting into (42) and using (43),

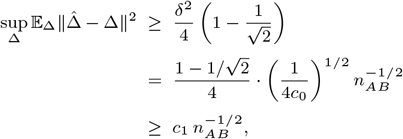

with 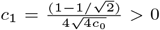. Since the left side is a lower bound on the minimax risk, this establishes (35). □

The rate 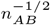 is slower than the usual parametric rate *n*^−1^ because Δ enters *p*_Δ_ only through the fourth cumulant of the arginal distribution. Although the heterozygous mixture has variance 1 + *δ*^2^*/*4 (differing from the null at second order), the *δ*^2^ perturbation to the log-density ratio is proportional to the second Hermite polynomial *H*_2_(*y*) = *y*^2^ − 1, which integrates to zero against the null density *p*_0_ by orthogonality. Consequently the leading distinguishing feature in the Chernoff exponent is a fourth-order quantity, so estimating Δ at the parametric rate yields squared error of order *n*^−1*/*2^. This is analogous to the rate for estimating the kurtosis of a distribution, and contrasts with mean or variance estimation (rate *n*^−1^).

When Σ_*AA*_ is unknown and estimated from *n*_*AA*_ homozygous observations, the whitening step introduces an additional error of order ∥Δ∥^2^ *· d/n*_*AA*_ (standard covariance estimation). The full minimax lower bound becomes

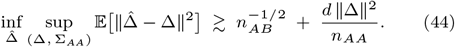

The first term is the irreducible cost of fourth-order estimation from heterozygotes; the second is the cost of learning the whitening transformation from homozygotes. Note however that given that *n*_*AA*_ ≫ *n*_*AB*_ and the scaling with *n*_*AA*_ as opposed to 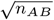 means even in a finite-sample regime estimation of the homozygous covariance is insignificant compared to estimation of the heterozygous largest eigenvalue/eigenvector.

### B.2. Numerical Evaluation

We then empirically compute how the Chernoff information between the standard Gaussian reference distribution *p*_0_ = *N* (0, *I*_*d*_) and 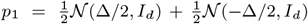 scales as a function of the separation magnitude ∥Δ∥. We evaluate this quantity on a logarithmically spaced grid of ∥Δ∥ values and uses a hybrid numerical strategy tailored to the two asymptotic regimes. For small ∥Δ∥, we use the leading-order analytic expansion *C*(*p*_0_, *p*_Δ_) ≈ ∥Δ∥^4^*/*256, which captures the quartic behavior near the origin. For larger ∥Δ∥, we use a Monte Carlo importance-sampling estimate based on the likelihood ratio *r*(*y*) = *p*_Δ_(*y*)*/p*_0_(*y*), compute the Chernoff objective 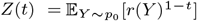, and numerically minimize − log *Z*(*t*) over *t* ∈ [0, 1]. The threshold between the analytic and Monte Carlo branches was chosen so that the Monte Carlo estimator is only used once it is numerically reliable.

After constructing this hybrid estimate, the we compare the resulting curve against two asymptotic descriptions. We plot the quartic reference curve ∥Δ∥^4^*/*256, which is expected to match the small-∥Δ∥ regime. Second, on the large-∥Δ∥ tail it fits a quadratic model of the form *a*∥Δ∥^2^ + *b*, testing the emergence of an approximately quadratic growth law at larger separations. The final output is a log-log plot showing the numerical Chernoff information together with these quartic and quadratic reference curves, with the small- and large-∥Δ∥ regions visually highlighted and the analytic-to-Monte-Carlo transition marked. The results are shown in Figure 9.

**Fig. 9:**
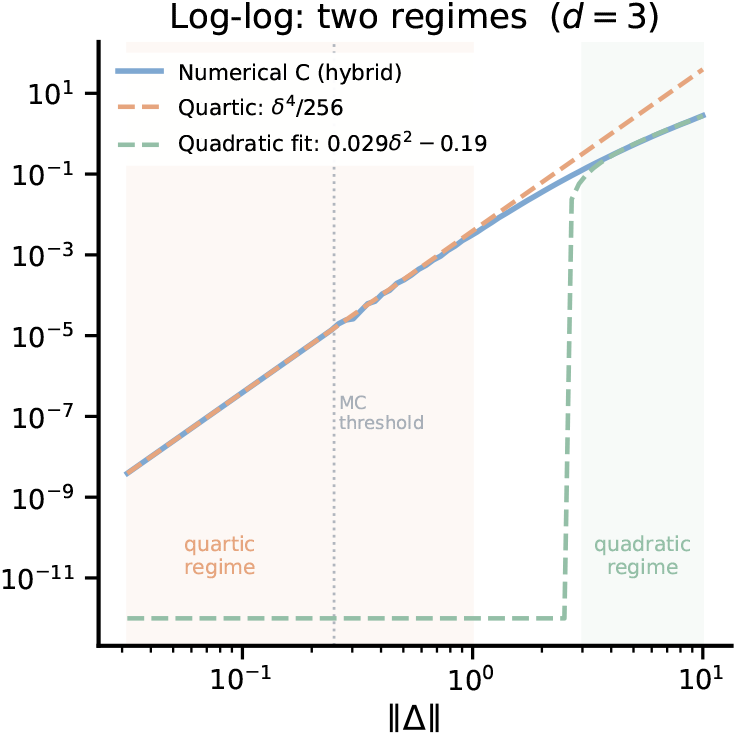
Scaling of Chernoff information between a standard Gaussian and a symmetric Gaussian mixture. Log-log plot of the Chernoff information *C*(*p*_0_, *p*_Δ_) between the reference distribution *p*_0_ = *N* (0, *I*_*d*_) and the mixture 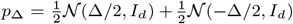, as a function of separation magnitude ∥Δ∥. The estimate is computed via a hybrid strategy: the analytic quartic approximation ∥Δ∥^4^*/*256 is used in the small-∥Δ∥ regime, while a Monte Carlo importance-sampling estimator is used for larger separations, with the transition point marked. Two asymptotic reference curves are overlaid: the quartic scaling ∥Δ∥^4^*/*256, which closely matches the small-separation behavior, and a fitted quadratic *a*∥Δ∥^2^ + *b*, which captures the emergent growth regime at large ∥Δ∥. Shaded regions highlight the two asymptotic regimes.

**Fig. 10:**
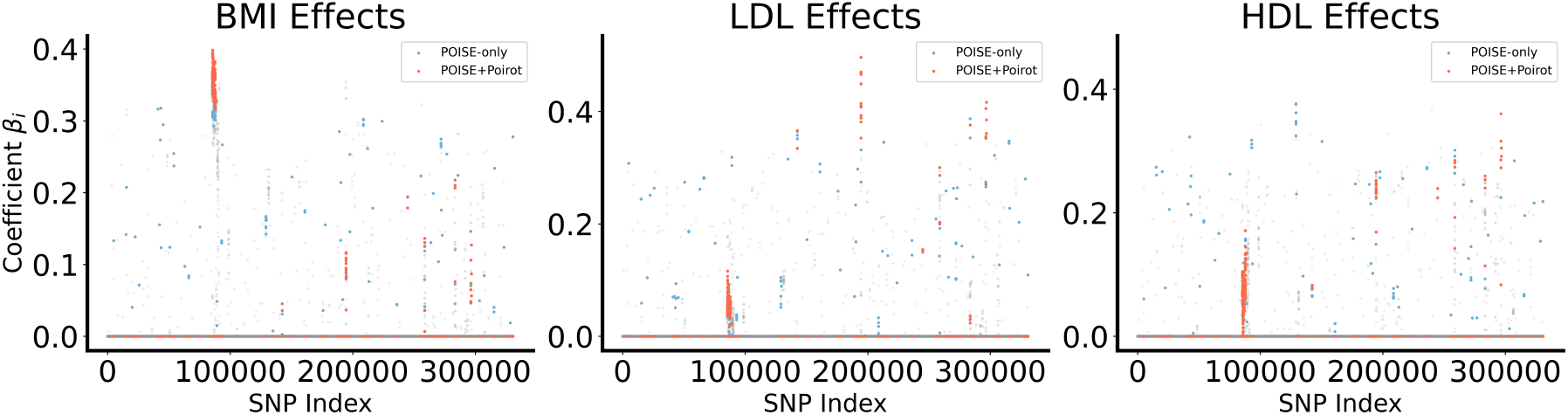
This figure matches Figure 6, but includes the remaining phenotypes, LDL and HDL.

## References

S. C. Barton, M. A. H. Surani, and M. L. Norris. Role of paternal and maternal genomes in mouse development. Nature, 311: 374–376, 1984.

A. Charcosset and A. Gallais. Estimation of the contribution of quantitative trait loci (qtl) to the variance of a quantitative trait by means of genetic markers. Theoretical and Applied Genetics, 93(8):1193–1201, 1996.

T. M. Cover and J. A. Thomas. Information theory and statistics. In Elements of Information Theory, chapter 11, pages 347–408. Wiley-Interscience, 2nd edition, 2006. ISBN 9780471241959.

B. Dumitrascu, G. Darnell, J. Ayroles, and B. E. Engelhardt. Statistical tests for detecting variance effects in quantitative trait studies. Bioinformatics, 35(2):200–210, 2019.

B. Efron. Better bootstrap confidence intervals. Journal of the American Statistical Association, 82(397):171–185, 1987. doi: 10.1080/01621459.1987.10478410.

B. Efron and R. J. Tibshirani. An introduction to the bootstrap. Chapman and Hall/CRC, 1994.

A. Guilmatre and A. Sharp. Parent of origin effects. Clinical genetics, 81(3):201–209, 2012.

S. T. Head, E. J. Leslie, D. J. Cutler, and M. P. Epstein. Poirot: a powerful test for parent-of-origin effects in unrelated samples leveraging multiple phenotypes. Bioinformatics, 39(4):btad199, 2023.

H. Hochner, C. Allard, E. Granot-Hershkovitz, J. Chen, C. M. Sitlani, S. Sazdovska, T. Lumley, B. McKnight, K. Rice, D. A. Enquobahrie, et al. Parent-of-origin effects of the apob gene on adiposity in young adults. PLoS genetics, 11(10):e1005573, 2015.

R. J. Hofmeister, S. Rubinacci, D. M. Ribeiro, A. Buil, Z. Kutalik, and O. Delaneau. Parent-of-origin inference for biobanks. Nature Communications, 13(1):6668, 2022.

R. J. Hofmeister, T. Cavinato, R. Karimi, A. van der Graaf, F.-D. Pajuste, J. Kronberg, N. Taba, E. B. research team Metspalu Andres 4 Esko Tõnu 4 Nelis Mari 4 Hudjashov Georgi 4, R. Mägi, M. Vaudel, et al. Parent-of-origin effects on complex traits in up to 236,781 individuals. Nature, 646(8085):647–656, 2025.

C. J. Hoggart, G. Venturini, M. Mangino, F. Gomez, G. Ascari, J. H. Zhao, A. Teumer, T. W. Winkler, N. Tšernikova, J. Luan, et al. Novel approach identifies snps in slc2a10 and kcnk9 with evidence for parent-of-origin effect on body mass index. PLoS genetics, 10(7):e1004508, 2014.

R. Howey, C. Mamasoula, A. Töpf, R. Nudel, J. A. Goodship, B. D. Keavney, and H. J. Cordell. Increased power for detection of parent-of-origin effects via the use of haplotype estimation. The American Journal of Human Genetics, 97(3):419–434, 2015.

S. J. Huxtable, P. J. Saker, L. Haddad, T. Love, R. P. Boot-Handford, H. Guassier, H. Brown, M. Walker, J. C. Levy, A. T. Hattersley, et al. Analysis of parent-offspring trios provides evidence for linkage and association between the insulin gene and type 2 diabetes mediated exclusively through paternally transmitted class iii variable number tandem repeat alleles. Diabetes, 49(1):126–130, 2000. doi: 10.2337/diabetes.49.1.126.

M. Lalande. Parental imprinting and human disease. Annual Review of Genetics, 30(1):173–195, 1996.

H. A. Lawson, J. M. Cheverud, and J. B. Wolf. Genomic imprinting and parent-of-origin effects on complex traits. Nature Reviews Genetics, 14(9):609–617, 2013.

L. LeCam. Convergence of estimates under dimensionality restrictions. The Annals of Statistics, pages 38–53, 1973.

O. Ledoit and M. Wolf. Honey, i shrunk the sample covariance matrix. The Journal of Portfolio Management, 30(4):110–119, 2004.

E. Li, C. Beard, and R. Jaenisch. Role for dna methylation in genomic imprinting. Nature, 366:362–365, 1993.

J. McGrath and D. Solter. Completion of mouse embryogenesis requires both the maternal and paternal genomes. Cell, 37: 179–183, 1984.

R. G. Miller. The jackknife-a review. Biometrika, 61(1):1–15, 1974.

R. D. Nicholls, J. H. M. Knoll, M. G. Butler, S. Karam, and M. Lalande. Genetic imprinting suggested by maternal heterodisomy in nondeletion prader-willi syndrome. Nature, 342:281–285, 1989.

W. Reik and J. Walter. Genomic imprinting: parental influence on the genome. Nature Reviews Genetics, 2(1):21–32, 2001.

C. Sudlow, J. Gallacher, N. Allen, V. Beral, P. Burton, J. Danesh, P. Downey, P. Elliott, J. Green, M. Landray, et al. Uk biobank: an open access resource for identifying the causes of a wide range of complex diseases of middle and old age. PLoS medicine, 12 (3):e1001779, 2015.

M. A. H. Surani, S. C. Barton, and M. L. Norris. Development of reconstituted mouse eggs suggests imprinting of the genome during gametogenesis. Nature, 308:548–550, 1984.

R. Vershynin. High-dimensional probability: An introduction with applications in data science, volume 47. Cambridge University press, 2018.

R. Weksberg, D. Ren Shen, Y. Ling Fei, Q. Li Song, and J. Squire. Disruption of insulin–like growth factor 2 imprinting in beckwith–wiedemann syndrome. Nature genetics, 5(2): 143–150, 1993.

B. Yu. Assouad, fano, and le cam. In Festschrift for Lucien Le Cam: research papers in probability and statistics, pages 423–435. Springer, 1997.

S. Zheng, Z. Bai, and J. Yao. Substitution principle for clt of linear spectral statistics of high-dimensional sample covariance matrices with applications to hypothesis testing. The Annals of Statistics, 43(2):546–591, 2015.

